# FOXO1 represses *Sprouty2* and *Sprouty4* expression in endothelial cells to promote arterial specification and vascular remodeling in the mouse yolk sac

**DOI:** 10.1101/2021.09.02.458792

**Authors:** Nanbing Li-Villarreal, Rebecca Lee Yean Wong, Monica D. Garcia, Ryan S. Udan, Ross A. Poché, Tara L. Rasmussen, Alexander M. Rhyner, Joshua D. Wythe, Mary E. Dickinson

**Affiliations:** Department of Molecular Physiology & Biophysics, Baylor College of Medicine. One Baylor Plaza, Houston, Texas 77030

## Abstract

The establishment of a functional circulatory system is required for post-implantation development during murine embryogenesis. Previous studies in loss of function mouse models have shown that FOXO1, a Forkhead family transcription factor, is required for yolk sac vascular remodeling and survival beyond embryonic day (E) 11. Here, we demonstrate that loss of *FoxO1* in E8.25 endothelial cells results in increased *Sprouty2* and *Sprouty4* transcripts, reduced expression of arterial genes, and decreased *Flk1/Vegfr2* mRNA levels without affecting overall endothelial cell identity, survival, or proliferation. Using a *Dll4-BAC-nlacZ* reporter line, we found that one of the earliest expressed arterial genes, *Delta like 4* (*Dll4)*, is significantly reduced in the yolk sac of *FoxO1* mutants without being substantially affected in the embryo proper. We show that in the yolk sac, FOXO1 not only binds directly to a subset of previously identified *Sprouty2* gene regulatory elements (GREs), as well as newly identified, evolutionarily conserved *Sprouty4* GREs, but can also repress their expression. Additionally, over expression of *Sprouty4* in transient transgenic embryos largely recapitulates reduced expression of arterial genes seen in endothelial *FoxO1* mutant mouse embryos. Together, these data reveal a novel role for FOXO1 as a key early transcriptional repressor controlling both pre-flow arterial specification and subsequent vessel remodeling within the murine yolk sac.

## INTRODUCTION

During early development the mammalian embryo requires a functional circulatory system to distribute oxygen, nutrients, and hormones. The mammalian heart is the first organ to form and function within this early embryo, along with the first arteries and veins that arise de novo via vasculogenesis (Fish and Wythe, 2015, Risau, 1994, Risau and Flamme, 1995, Chong et al., 2011). Primitive erythrocytes form in the blood islands of the extra-embryonic yolk sac (YS), and are drawn into circulation as the heart begins to beat around embryonic day (E) 8 in the mouse embryo (Lucitti et al., 2007, Palis, 2014, Ji et al., 2003). A complete circulatory loop between the embryo and the extra-embryonic YS is evident shortly after the onset of cardiac contractions, with blood flowing through the dorsal aorta to the vitelline (omphalomesenteric) artery (VA), through the YS capillary plexus, and back through the vitelline (omphalomesenteric) vein (VV) to the sinus venosus of the heart. Circulation through the YS is the main circulatory loop until the chorion-allantoic placenta connections develop later around E9.

A key finding several years ago showed that arterial-venous (AV) identity is established prior to the onset of blood flow in the early mouse embryo (Herzog et al., 2005, Chong et al., 2011, Aitsebaomo et al., 2008, Wang et al., 1998, Adams et al., 1999). Others have since shown that some aspects of AV identity are plastic, as they can be influenced by changes in blood flow and hemodynamics (le Noble et al., 2004, le Noble et al., 2005, Wragg et al., 2014). Extensive work in zebrafish has shown that AV specification depends on differential responses to VEGF signaling through the tyrosine kinase receptor VEGFR2/Flk1, high levels of which activate the MEK/ERK kinase cascade in arterial cells and PI3K/AKT signaling in venous cells (Fish and Wythe, 2015, Covassin et al., 2006, Weinstein and Lawson, 2002). In the arterial endothelium, VEGFR2 signaling stimulates the Notch pathway, which in turn promotes an arterial identity while simultaneously repressing a venous fate (Weinstein and Lawson, 2002, Krebs et al., 2010, Lawson et al., 2001, Lawson et al., 2002, Liu et al., 2003, Shutter et al., 2000, Swift and Weinstein, 2009, Siekmann and Lawson, 2007, Krebs et al., 2000, Duarte et al., 2004, Lobov et al., 2007). VEGF upregulates expression of *Delta-like 4 (Dll4)*, which encodes a ligand for the Notch family of transmembrane receptors. *Dll4* is the earliest Notch ligand expressed in arterial cells in the early mouse embryo (Shutter et al., 2000, Mailhos et al., 2001, Wythe et al., 2013, Cleaver and Krieg, 1998, Chong et al., 2011), and is essential for AV patterning (Krebs et al., 2000, Gale et al., 2004, Duarte et al., 2004). Expression of *Dll4* depends on the activation of ETS transcription factors, downstream of VEGF signaling (Wythe et al., 2013, Fish et al., 2017) and *Dll4* mRNA expression can be increased by shear stress (Masumura et al., 2009, Obi et al., 2009). However, the expression of *Dll4* and a few other select arterial markers prior to, and independent of, the onset of blood flow (Chong et al., 2011, Wang et al., 1998) suggests flow-independent mechanisms regulate arterial specification. Vegfr2 also upregulates expression of the main endothelial cell surface receptor for Dll4, Notch1 (Lawson et al., 2002). Notch itself is activated by blood flow and the strength of this signal depends on the magnitude of shear stress (Masumura et al., 2009, Mack et al., 2017). Notch regulates cell junctions, cell cycle arrest, and induces an arterial gene expression program via a connexin37 (*Gja4*)/p27^Kip1^ (*Cdkn1B*) pathway (Mack et al., 2017, Su et al., 2018, Fang et al., 2017). Critically, both Vegfr2 and Notch1 are thought to act as mechanosensors, likely linking arterial specification of ECs and hemodynamic feedback via blood flow to re-enforce and solidify their arterial identity (Mack et al., 2017, Tzima et al., 2005, Mack and Iruela-Arispe, 2018, Shay-Salit et al., 2002).

Forces exerted by blood flow play a clear role in AV specification, and also influence vessel morphogenesis and remodeling in the early embryo (Fang et al., 2017, Masumura et al., 2009, le Noble et al., 2004, Chong et al., 2011, Hwa et al., 2017). Hemodynamic force is both necessary and sufficient to remodel the high-resistance mouse YS capillary plexus into a more complex hierarchical network with a large caliber VA and VV progressively leading to smaller diameter vessels (Lucitti et al., 2007). Our lab has shown that murine yolk sac vessel remodeling depends on both vessel fusion and EC migration (Udan et al., 2013). Interestingly, live imaging studies showed distinct differences in how arterial and venous cells respond to changes in hemodynamic force (Udan et al., 2013, Kondrychyn et al., 2020, Goetz et al., 2014). These data suggest that pathways that control AV identity, which can be regulated by blood flow, may also influence physical responses of ECs to blood flow such as migration and motility that facilitate vessel remodeling.

Despite the knowledge that has been gained regarding the mechanisms regulating arterial-venous identity and the discovery of mechanosensors that are required for ECs to sense blood flow, a full understanding of these mechanistic pathways is yet to be realized. A recent analysis estimates that approximately 6% of genes in the genome (∼1200) may be required during early cardiovascular development (E9.5-E12.5) (Dickinson et al., 2016). One such gene, and the focus of this study, is *FoxO1 (Forkhead box protein O1). Forkhead domain class O transcription factors* (*FOXOs)* integrate different cellular signaling pathways to regulate cellular homeostasis (Paik et al., 2007, Huang and Tindall, 2007, Jiramongkol and Lam, 2020). *Daf-16/FoxO* was originally identified as a regulator of dauer formation in *C. elegans* (Albert et al., 1981) and was later shown to control longevity by sensing environmental cues such as hormones, nutrient availability, oxidative stress, and energy metabolism via signaling through the insulin, AKT/mTor, JNK and AMPK pathways (Sun et al., 2017). Several studies have since established that *FoxO1* is also required for normal embryonic development in mice. Homozygous null *FoxO1* embryos display a primitive yolk sac vasculature, pericardial edema, and disorganized embryonic vessels by E9.5, resulting in lethality by E11.5 (Furuyama et al., 2004, Dharaneeswaran et al., 2014, Hosaka et al., 2004, Sengupta et al., 2012, Wilhelm et al., 2016, Ferdous et al., 2011). Further analysis showed that loss of *FoxO1* in the endothelium, but not the myocardium, phenocopied germline loss of *FoxO1* (Sengupta et al., 2012), demonstrating the cell autonomous requirement for FOXO1 in the embryonic vasculature. Follow-up studies have since found that FOXO1 controls a variety of different processes in endothelial cells (ECs), including, but not limited to EC proliferation and metabolism (Wilhelm et al., 2016), endothelial barrier function (Beard et al., 2020), sprouting angiogenesis (Kim et al., 2019, Fukumoto et al., 2018, Dang et al., 2017), autophagy (Zhang et al., 2019), EC growth (Riddell et al., 2018, Rudnicki et al., 2018) and migration (Niimi et al., 2017). Despite these studies, the exact function that FOXO1 plays in the early vasculature remains elusive. Given that FOXO1 activity can be modulated in response to fluid shear stress (Chlench et al., 2007, Dixit et al., 2008), combined with our studies showing that hemodynamic force is necessary and sufficient for early vascular remodeling (Lucitti et al., 2007, Udan et al., 2013), and the growing evidence for FOXO1 in cell migration and sprouting angiogenesis (Fosbrink et al., 2006, Niimi et al., 2017, Kim et al., 2019), we define the requirement for FOXO1 in the remodeling vasculature of the early embryonic yolk sac.

Herein, we demonstrate a novel role for FOXO1 in regulating AV identity in the murine YS vasculature. Using conditional, endothelial-specific *FoxO1* loss-of-function mutants, we identified a significant down regulation of arterial gene expression in the mouse yolk sac prior to the onset of blood flow. We also detected a significant reduction in *Vegfr2/Flk1* transcripts, but normal expression levels for other pan-endothelial genes such as *Pecam1*, indicating that the formation of ECs is not disrupted but rather VEGF signaling is affected. Using a novel *Dll4* arterial reporter line, we showed that *FoxO1* is required for *Dll4* expression in the murine yolk sac, but not in the embryo proper. Further analysis showed that FOXO1 represses expression of *Sprouty* genes, which encode inhibitors of Raf/MEK/ERK signaling downstream of FGF and VEGF receptor activation. Sprouty factors also modulate angiogenesis by negatively regulating small vessel branching, as well as repressing endothelial cell migration (Gong et al., 2013, Wietecha et al., 2011, Lee et al., 2001). While some have shown that FOXO1 positively regulates *Sprouty* gene expression (Paik et al., 2007), our studies demonstrate that *FoxO1* loss increased *Sprouty2* and *4* mRNA levels, suggesting that FOXO1 represses *Sprouty2*/*4* in the murine yolk sac. We went on to find that *Sprouty4* overexpression throughout the yolk sac and embryo profoundly altered arterial gene expression in the yolk sac but, similar to early FoxO1 loss, had an insignificant effect on these transcripts in the embryo proper. Taken together, these data highlight a novel role for FOXO1 in regulating arteriovenous specification in the early yolk sac and reveal a new mechanism wherein FOXO1 represses *Sprouty* gene expression and downstream signaling in the endothelium.

## Results

### Defective yolk sac vascular remodeling in *FoxO1*^ECKO^ embryos is not due to abnormalities in hemodynamic force or allantois defects

To define the role of FOXO1 in ECs within the early embryo, we conditionally ablated *FoxO1* in the endothelium by crossing *FoxO1*^*flox*^ mice (Paik et al., 2007) with *Tie2-Cre* transgenic mice, in which Cre recombinase is expressed in endothelial and hematopoietic cells and progenitors starting at E7.5 (Kisanuki et al., 2001). The efficiency of the *Tie2-Cre* mediated recombination of the *FoxO1*^flox^ allele was confirmed by qRT-PCR, which showed over 60% reduction in *FoxO1* mRNA in CKO yolk sacs compared to control littermates at E8.5 (Figure 1A). We observed gross phenotypes similar to previous reports (Figure 1B and C and S1A and B) (Sengupta et al., 2012, Furuyama et al., 2004, Hosaka et al., 2004). There were no visible differences in vascular morphology or embryo size between control (*Tie2-Cre*^*+/+*^*;FoxO*^*+/flox*^) and <u>e</u>ndothelial <u>c</u>onditional <u>k</u>nock<u>o</u>ut embryos (*Tie2-Cre*^*+/tg*^*;FoxO1*^*flox/flox*^, hereafter referred to as *FoxO1*^*ECKO*^) at E8.5 (Figures 1D and E). However, at E9.5 and E10.5, while the primitive vascular plexus of the control yolk sac had remodeled into a hierarchy of large caliber vessels iteratively branching into smaller diameter capillaries, *FoxO1*^*ECKO*^ yolk sacs retained a primitive vascular plexus (Figures 1F-I). *FoxO1*^*ECKO*^ embryos stained with anti-CD31 antibodies showed a thinner and less branched vitelline vein and artery compared to controls (Figure 1J). At E9.5, mutant embryos were reduced in overall size compared to control littermates (Figures 1F and G), and this became more evident at E10.5 (Figures 1B-C, H-I). Additionally, in both E9.5 (not shown) and E10.5 *FoxO1*^*ECKO*^ embryos (Figure 1I), the pericardial sac was enlarged, and blood was abnormally pooled in the heart. Overall, phenotypes in *FoxO1* mutant embryos are highly reproducible and our observations align well with previously published studies (Sengupta et al., 2012, Furuyama et al., 2004, Hosaka et al., 2004).

**Figure 1.**
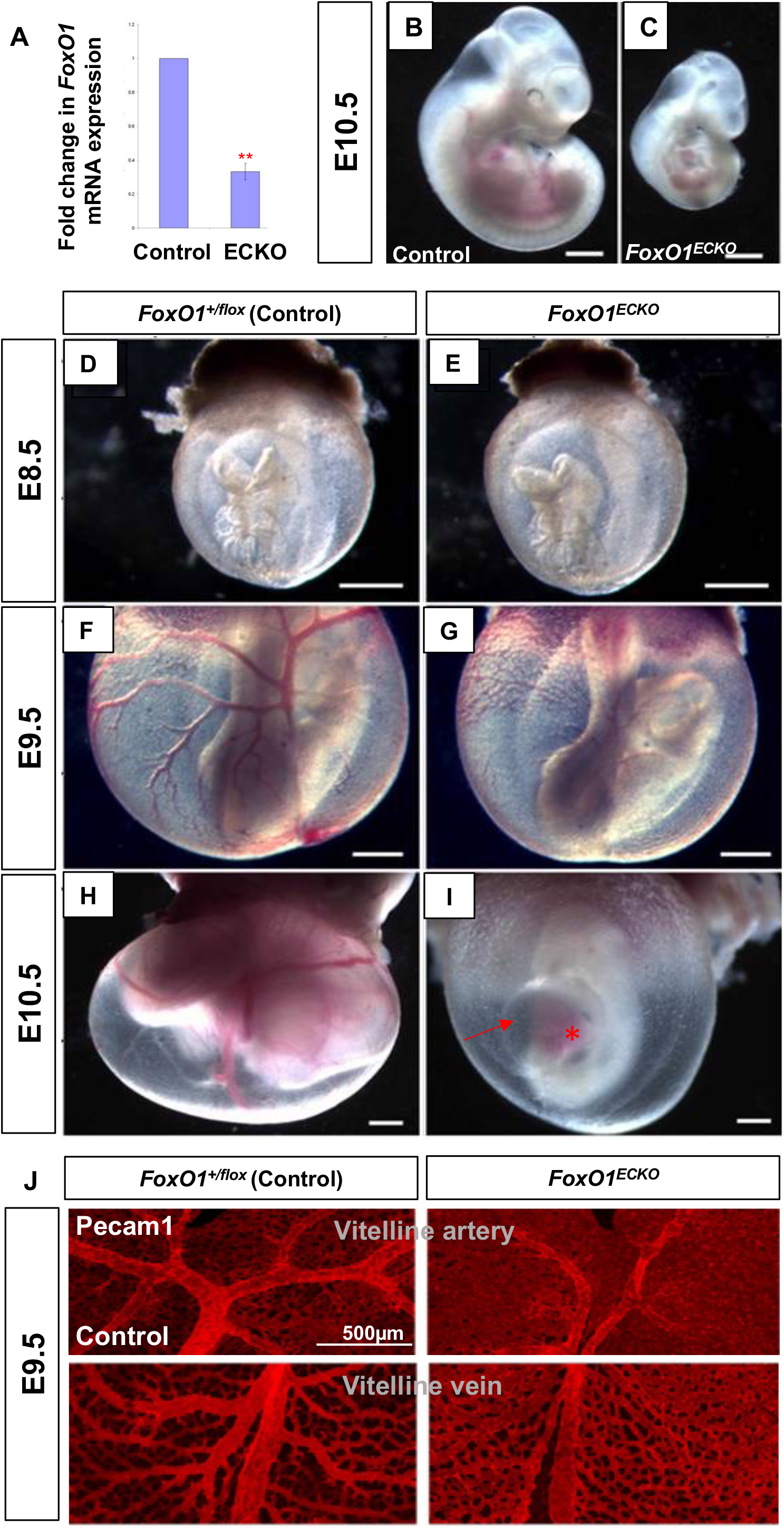
*FoxO1*^*ECKO*^ results in vascular remodeling defects and lethality. (A) Quantitative RT-PCR for *FoxO1* expression in control and ECKO yolk sacs. *P*<0.01. (B and C) Bright field images of E10.5 littermate control and *FoxO1*^*ECKO*^ embryos. Control and *FoxO1*^*ECKO*^ embryos within the yolk sac at E8.5 (D and E), E9.5 (F and G), and E10.5 (H and I). Pericardial edema (arrow), blood pooling in the heart (asterisk). (J) Pecam1 staining in E9.5 control and CKO yolk sacs. Scale bar = 500μm.

The appearance of pericardial edema in *FoxO1*^*ECKO*^ embryos is suggestive of heart failure and compromised circulation. To determine if and when blood flow was impaired, we crossed a primitive erythrocyte transgenic fluorescent reporter line, *ɛ-globin-KGFP* (Dyer et al., 2001) into the *FoxO1*^*ECKO*^ background. Live imaging of cultured embryos and high-speed confocal microscopy was used to track individual KGFP labeled erythroblasts to determine blood velocity (Figure 2) (Jones et al., 2004). At E8.5, both control and *FoxO1*^*ECKO*^ embryonic vessels were filled with blood and erythrocytes moved with a steady directional flow with similar periodicity and velocity (Figures 2A, B, E, and G). By E9.5, control embryos had clearly remodeled vessels, blood flow velocity greater than 700 μm/s, and a defined wave pattern with periodicity of 400 ms (Figures 2C and F). However, *FoxO1*^*ECKO*^ embryos vessels were not remodeled, and blood flow had a significantly lower velocity, with a poorly defined wave pattern (Figures 2D and H). Quantification of blood velocity (Figures 2I-J) and heart rate (Figures 2K-L) revealed no statistical difference in E8.5 control and *FoxO1*^*ECKO*^ embryos (Figures 2I and K). However, by E9.5, blood velocity and heart rate were significantly decreased in *FoxO1*^*ECKO*^ embryos (Figures 2J and L), indicating heart failure was occurring in embryos with un-remodeled vessels. Thus, flow initiates normally in *FoxO1*^*ECKO*^ embryos and flow abnormalities are not detected until after defects in vessel remodeling are evident. Given these results, we restricted our analysis, whenever possible, to E8.25-E8.5 embryos so that poor blood flow did not influence our observations.

**Figure 2.**
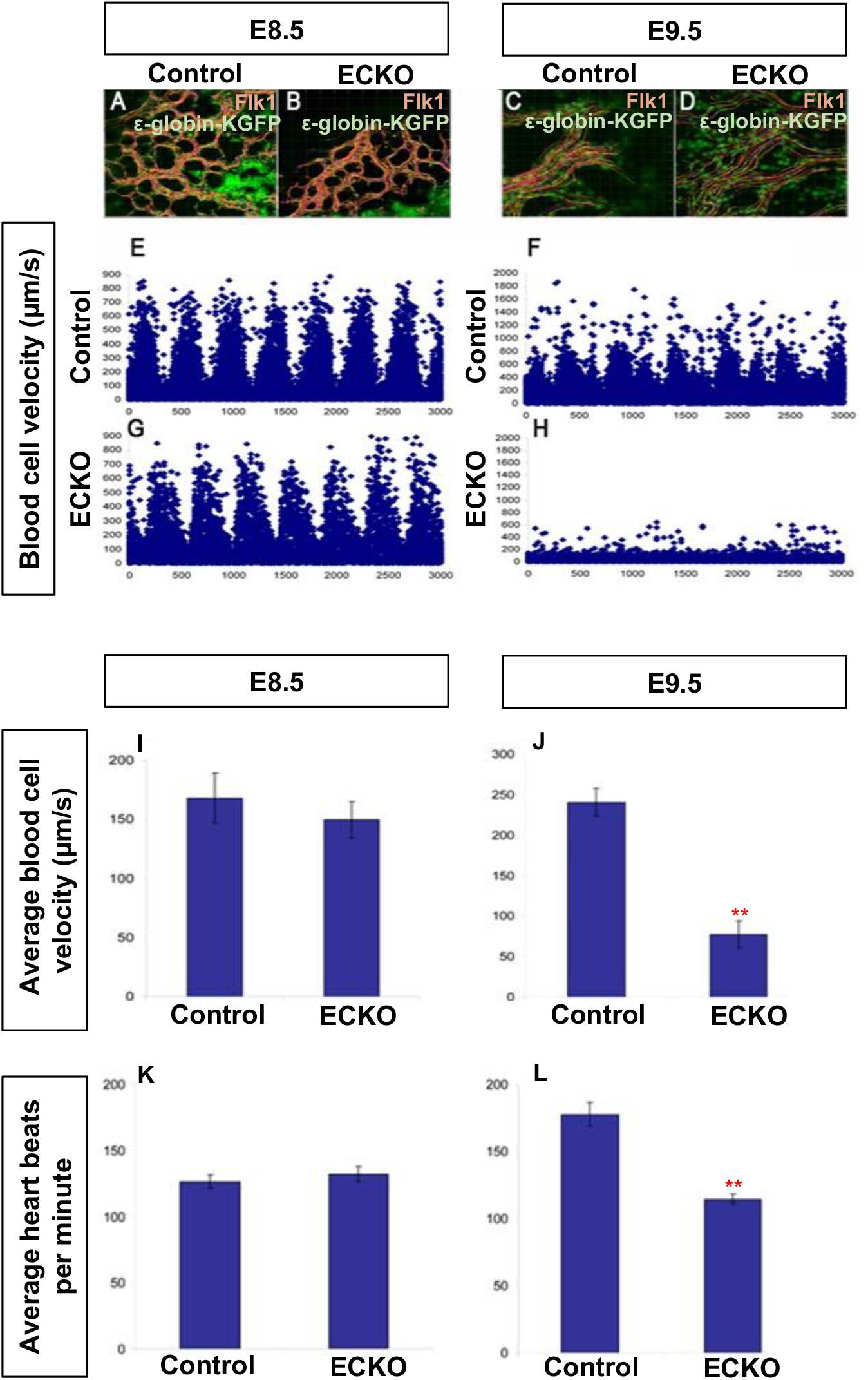
Vascular remodeling defects in *FoxO1*^*ECKO*^ embryos do not result from reduced blood flow. Primitive erythroblasts in circulation in wild type and CKO embryos were marked by crossing to an *ε-globin-GFP* transgenic reporter. Representative still images of E8.5 wild type (A), ECKO (B), E9.5 wild type (C), and ECKO (D) embryos. Individual blood cells from A-D were tracked and velocity profiles are plotted in E-H. Quantification of the average blood velocity are graphed in I and J (Mann-Whitney U test, p=0.005). Average heart rates quantified in wild type and ECKO embryos at E8.5 (K) and E9.5 (L) (Kruskal-Wallis test, p=0.003). Bars in graphs are means ± standard error.

A previous report on the role of *FoxO1* in placental development described phenotypes including swollen or hydropic allantois, failed chorion-allantoic fusion, and increased cell death in the allantois (Ferdous et al., 2011). Since the previous study was conducted in germline *FoxO1* null embryos, we examined the allantois in global null and *FoxO1*^*ECKO*^ mutants. While germline *FoxO1* mutants exhibited partially penetrant defects in allantois formation and fusion, these phenotypes were not evident in *FoxO1*^ECKO^ embryos at E9.5 (Table 1). However, both germline and *FoxO1*^*ECKO*^ embryos show defects in yolk sac vascular remodeling, heart failure phenotypes, and lethality by E11.5. Taken together, these results support a cell autonomous requirement for FOXO1 in yolk sac vessel remodeling and suggest that published cardiac and blood flow defects are secondary to the impaired remodeling of yolk sac vasculature.

**Table 1.** Allantois phenotype analysis in control, null and ECKO embryos at E9.5.

|  | Genotype | Number | Percent of Total | Gross Allantois Defects | Percent of genotype |
| --- | --- | --- | --- | --- | --- |
| 4 Litters | WT | 16 | 31.37 |  |  |
|  | *FoxO1<sup>floxdel/+</sup>* | 20 | 39.21 |  |  |
|  | *FoxO1<sup>floxdel/floxdel</sup>* | 15 | 29.41 | 2 | 13.33 |
|  | Total | 51 |  |  |  |
| 6 Litters | *FoxO1<sup>flox/+</sup>* | 15 | 31.91 |  |  |
|  | *FoxO1<sup>flox/+</sup>;Tie2-cre<sup>Tg/+</sup>* | 12 | 25.53 |  |  |
|  | *FoxO1<sup>flox/flox</sup>* | 10 | 21.27 |  |  |
|  | *FoxO1<sup>flox/flox</sup>;Tie2-cre<sup>Tg/+</sup>* | 10 | 21.27 | 0 | 0 |
|  | Total | 47 |  |  |  |

### FOXO1 is necessary to maintain *Flk1/Vegfr2* expression in E8.25 embryos

To further assess the effect of endothelial-specific *FoxO1* deletion in the developing embryo, we examined transcript levels of genes normally expressed in blood vessels by qRT-PCR within *FoxO1*^*ECKO*^ E8.25 yolk sacs. We focused these experiments at E8.25 to avoid potential complications of later stage heart failure. We observed a significant decrease in *Flk1/Vegfr2* expression in *FoxO1*^*ECKO*^ yolk sacs, while other pan-endothelial markers such as *Pecam1, Tie2, VE-Cadherin, Flt1/Vegfr1*, and *Cx43* were not significantly affected (Figure 3A). Immuno-labeling of endogenous *Flk1* showed a similar reduction in expression in *FoxO1*^*ECKO*^ yolk sac endothelial cells at E8.25, while Pecam1 (CD31) expression – a pan endothelial cell marker – appeared unaffected (Figure 3B). We further confirmed the reduction in *Flk1* expression using magnetic-activated cell sorting (MACS) to isolate CD31^+^ ECs from E8.25 WT and *FoxO1* germline deletion mutant yolk sacs. CD31^+^ yolk sac ECs showed seven-fold higher expression of *Pecam1* mRNA (which encodes CD31) than whole yolk sacs (isolated separately then combined) and approximately thirty-fold higher than the non-endothelial population (CD31^-^), demonstrating the enrichment of endothelial cells via MACS (Figure 3C). To compare WT and mutant endothelial cells, we assayed transcript levels of both *Pecam1* (CD31) and *Flk1* (Vegfr2) in the CD31^+^ populations. *Pecam1* levels were comparable between WT and mutant endothelial cells, whereas *Flk1* was significantly reduced in the germline mutant endothelial cells (Figure 3D and E). Finally, we examined *Flk1* expression using a transgenic reporter, *Flk1-H2B::YFP* (Fraser et al., 2005), which labels endothelial cell nuclei. At E8.25, YFP^+^ endothelial nuclei were evenly dispersed throughout the vascular plexus of control yolk sacs, however, the number of YFP^+^ nuclei within the *FoxO1*^*ECKO*^ yolk sacs were significantly reduced (Figure 3F). The total number of DAPI^+^ nuclei was unchanged. Nuclear segmentation and quantification of the average YFP^+^ cell density revealed a significant reduction in the number of YFP^+^ cells in *FoxO1*^*ECKO*^ yolk sacs compared to controls (Figure 3G). To determine if the reduction in YFP^+^ cells was due to a difference in apoptosis or proliferation, *FoxO1*^*ECKO*^ litters positive for the *Flk1-H2B-YFP* reporter were immunostained for phospho-histone 3 (PH3) or Caspase 3 (Figure 3H and I). We found that neither cell proliferation nor apoptosis within the yolk sac differed significantly between control and *FoxO1*^*ECKO*^ embryos (Figures 3 H, I, S2A-D). Since the mRNA expression of pan-endothelial markers was not decreased, but Flk1 transcripts and protein were reduced, we concluded that the low density in YFP^+^ cells is not due to reduced EC numbers, but rather reduced expression of the *Flk1* reporter. Collectively, these data indicate that FOXO1 is required to maintain cell autonomous *Flk1/Vegfr2* expression in ECs, but not required for the formation, proliferation or survival of ECs or the expression of other pan-endothelial markers.

**Figure 3.**
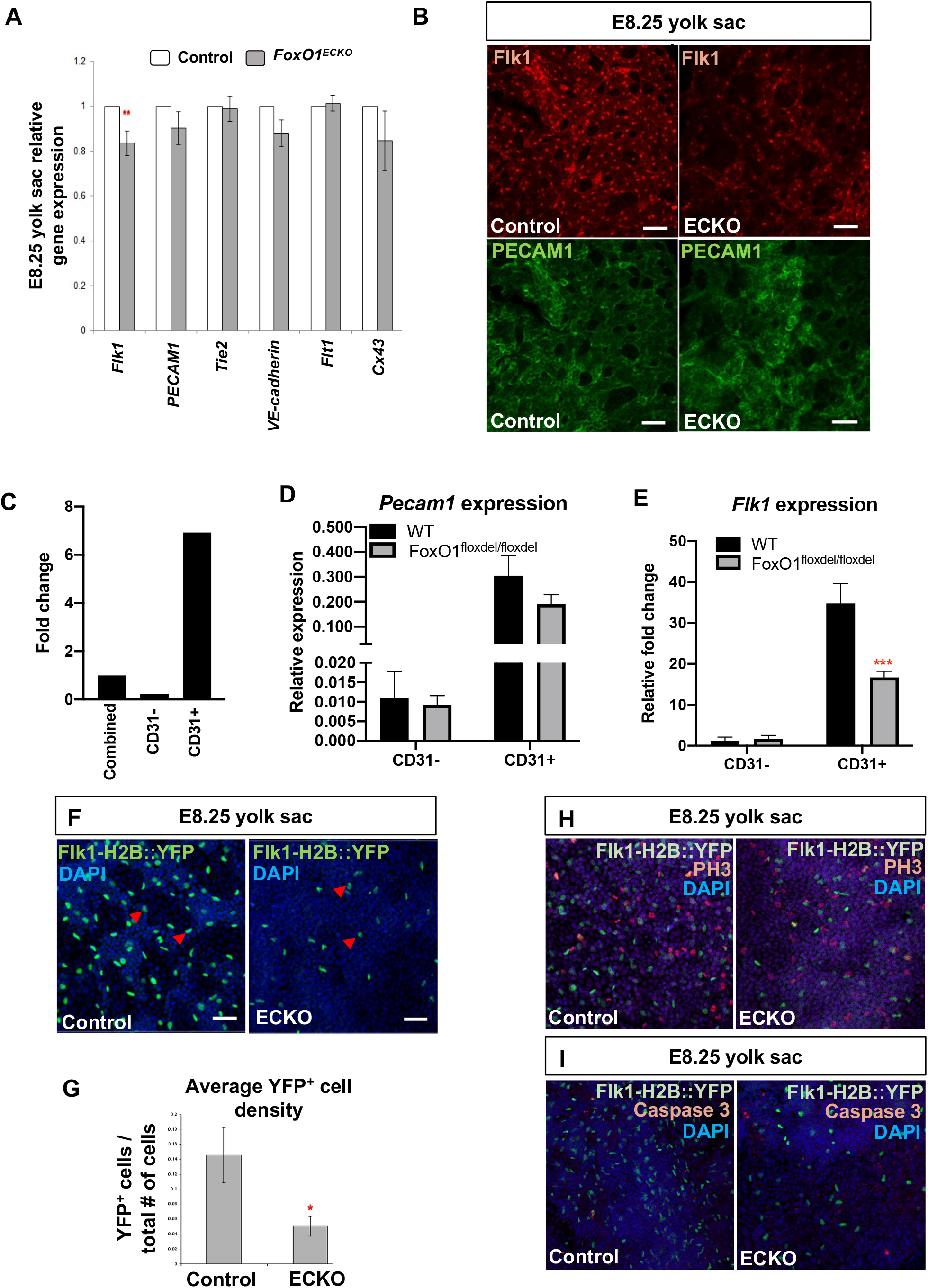
FOXO1 regulates FLK1 expression without affecting other endothelial genes or endothelial cell viability prior to blood flow. (A) Expression levels of endothelial genes by quantitative RT-PCR. (B)Yolk sacs from control and *FoxO1*^*ECKO*^ at E8.25 were DAPI-stained and labelled with *Flk1-H2B::YFP* transgene, which marks the nuclei of endothelial cells (arrowheads). Whole mount phosphoHistone-H3 (PH3) (C) or activated Caspase3 (D) staining of control and *FoxO1*^*ECKO*^ E8.25 yolk sacs co-labeled with *Flk1-H2B::YFP* transgene and DAPI. (E) Immuno-labeling for endogenous Flk1 and Pecam1 in control and *FoxO1*^*ECKO*^ yolk sacs. Scale bars = 50μm (B -E). (F) Comparison of *Pecam1* expression in MACS sorted CD31-, CD31+, and combined control E8.25 yolk sac cells by qPCR. Relative *Pecam1* (G) and *Flk1* (H) expression between WT and *FoxO1* null E8.25 MACS sorted CD31-and CD31+ yolk sac cells.

### FOXO1 is required to regulate arterial gene expression in the yolk sac vasculature

Given the reduction in *Flk1/Vegfr2* expression in E8.25 embryos, and the critical role of VEGF-VEGFR2 signaling in establishing arteriovenous identity in the early embryonic endothelium, we next examined other markers of AV specification (Figure 4A). Previously, Furuyama *et al*. showed reduced expression of the arterial-enriched transcripts *Cx40, Cx37, eNOS, and EphrinB2* in the yolk sacs of E9.5 *FoxO1*^*ECKO*^ mutants compared to control littermates (Furuyama et al., 2004), but given the changes in blood flow that we observed in E9.5 *FoxO1*^*ECKO*^ embryos, we were interested to determine if expression of these markers was affected earlier. Indeed, these genes were significantly reduced in *FoxO1*^*ECKO*^ yolk sacs at E8.25 compared to controls (Figure 4A). In addition, we found a significant reduction in transcript levels of Notch family members, including *Notch1, Hey1, Hey2, Jagged1* and *Dll4*, which are required for arterial specification (Gridley, 2010, Duarte et al., 2004, Xue et al., 1999, Fischer et al., 2004). Furthermore, *Neuropilin1*, which encodes a co-receptor for VEGFR2 and is required for arterial specification, was also downregulated, whereas its venous counterpart, *Neuropilin2*, remained unchanged (Herzog et al., 2001). Additional venous markers, *Coup-TFII (NR2F2)* and *EphB4*, showed a modest decrease, or no significant change, respectively (Wang et al., 1998, You et al., 2005). The endodermal marker *Afp* was also unchanged (Figure 4A) (Dziadek and Adamson, 1978), showing further that gene transcription changes were confined to markers in the arterial endothelium. These results demonstrate that FOXO1 is required for normal expression of early arterial genes, but is dispensable for venous identity, in the murine yolk sac.

**Figure 4.**
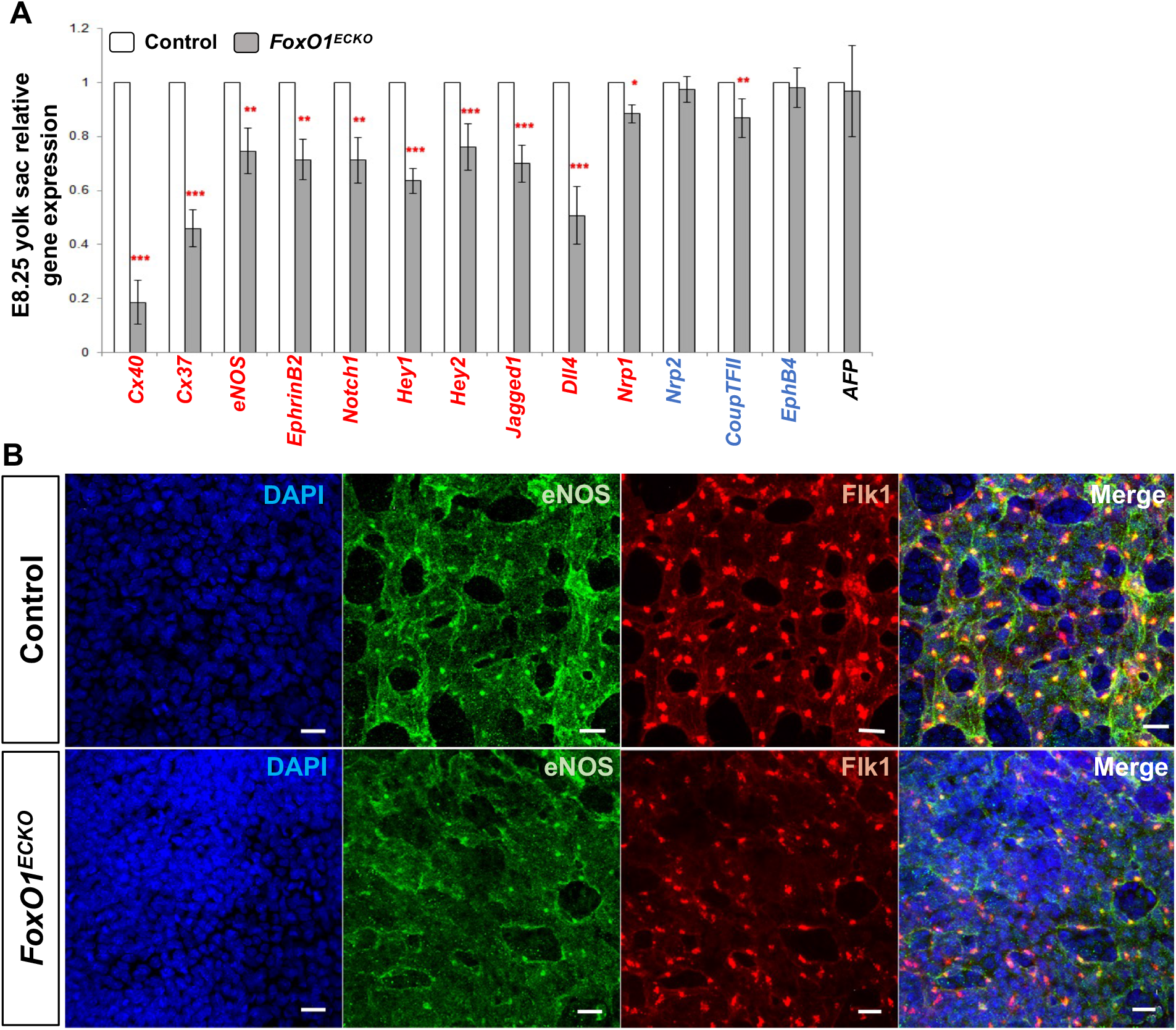
Arterial marker expression is reduced in *FoxO1*^*E*CKO^ yolk sac. (A) Quantitative RT-PCR expression analysis in control and *FoxO1*^*ECKO*^ yolk sacs; arterial (red), venous (blue), and endoderm markers. \**p*<0.05, \*\**p*<0.01, \*\*\**p*<0.001, data are means ±S.E. (B) Immuno-labeling of eNOS and FLK1 in control and *FoxO1*^*ECKO*^ yolk sacs. Scale bars = 20μm.

To determine if reduced transcript levels of arterial-specific genes correlated with decreased expression of their respective proteins in *FoxO1*^*ECKO*^ yolk sacs, immunofluorescence was performed on sectioned yolk sacs at E8.25. Confocal imaging of Cx37 and Cx40 revealed an overall reduction in the number of connexin-positive puncta in *FoxO1*^*ECKO*^ yolk sacs when compared to controls (Figure S3A and B). Similarly, we observed decreased eNOS expression within the vascular plexus in *FoxO1*^*ECKO*^ yolks sacs compared to controls (Figure 4B). These data, in addition to previous gene expression analysis, indicate that FOXO1 within the developing endothelium is necessary for the regulation of arterial endothelial cell identity.

### Characterization of arterial defects in *FoxO1*^*ECKO*^ and germline mutants using the *Dll4-BAC-nlacZ* reporter

Thus far, our phenotypic, transcriptional, and immunolabeling studies support a role for FOXO1 as a regulator of vascular remodeling, *Flk1* expression and arterial specification of endothelial cells within the yolk sac. To further analyze the arterial specification defects in *FoxO1*^*ECKO*^ mutants, we examined the spatial expression of one of the earliest markers of arterial identity (Chong et al., 2011, Wythe et al., 2013), *Dll4*, using a transgenic reporter line, *Dll4-BAC-nlacZ*, that faithfully recapitulates endogenous *Dll4* expression (Herman et al., 2018). E8.25 *FoxO1*^*ECKO*^ embryos carrying the nuclear-localized β-galactosidase reporter were compared with control littermates (Figure 5). In E8.25 control embryos, *Dll4-BAC-nlacZ* reporter activity was observed in the dorsal aorta (DA), endocardium (EC), and nascent umbilical artery (UA) within the allantois (Figure 5A and S4A). LacZ positive nuclei were also detected within the YS in and around the vitelline artery and arterioles (Figure 5A, red arrows). *FoxO1*^*ECKO*^ yolk sacs exhibited a similar LacZ expression pattern spatially, albeit with reduced intensity (Figure 5B), consistent with our transcript analysis showing reduced *Dll4* mRNA expression in the *FoxO1*^*ECKO*^ yolk sacs (Figure 4A and 5C). However, unlike in the yolk sac, nLacZ expression was only slightly reduced in the embryo proper of *FoxO1*^*ECKO*^ mutants (Figure S4B compared to S4A, Figure 5C). To determine if the differences in nLacZ reporter expression between the yolk sac and embryo was influenced by Cre-mediated recombination in our conditional knockout studies, we examined the activity of the *Dll4* LacZ reporter in germline *FoxO1* mutants. Figures 4SE and F show representative images of anterior views of the embryos, while Figure 5D and E show posterior views. Results similar to the findings in the *FoxO1*^*ECKO*^ embryos were observed, but with even greater decreases in reporter expression between control (5D red arrows) and null (5E red arrows) yolk sacs and embryos. Endogenous *Dll4* expression in the embryo proper and the yolk sac showed a small, but significant decrease in endogenous *Dll4* expression between control and *FoxO1* null embryos. Comparatively, a dramatic reduction in *Dll4* expression was observed in the *FoxO1* null yolk sacs compared to wildtype controls (Figure 5F).

**Figure 5.**
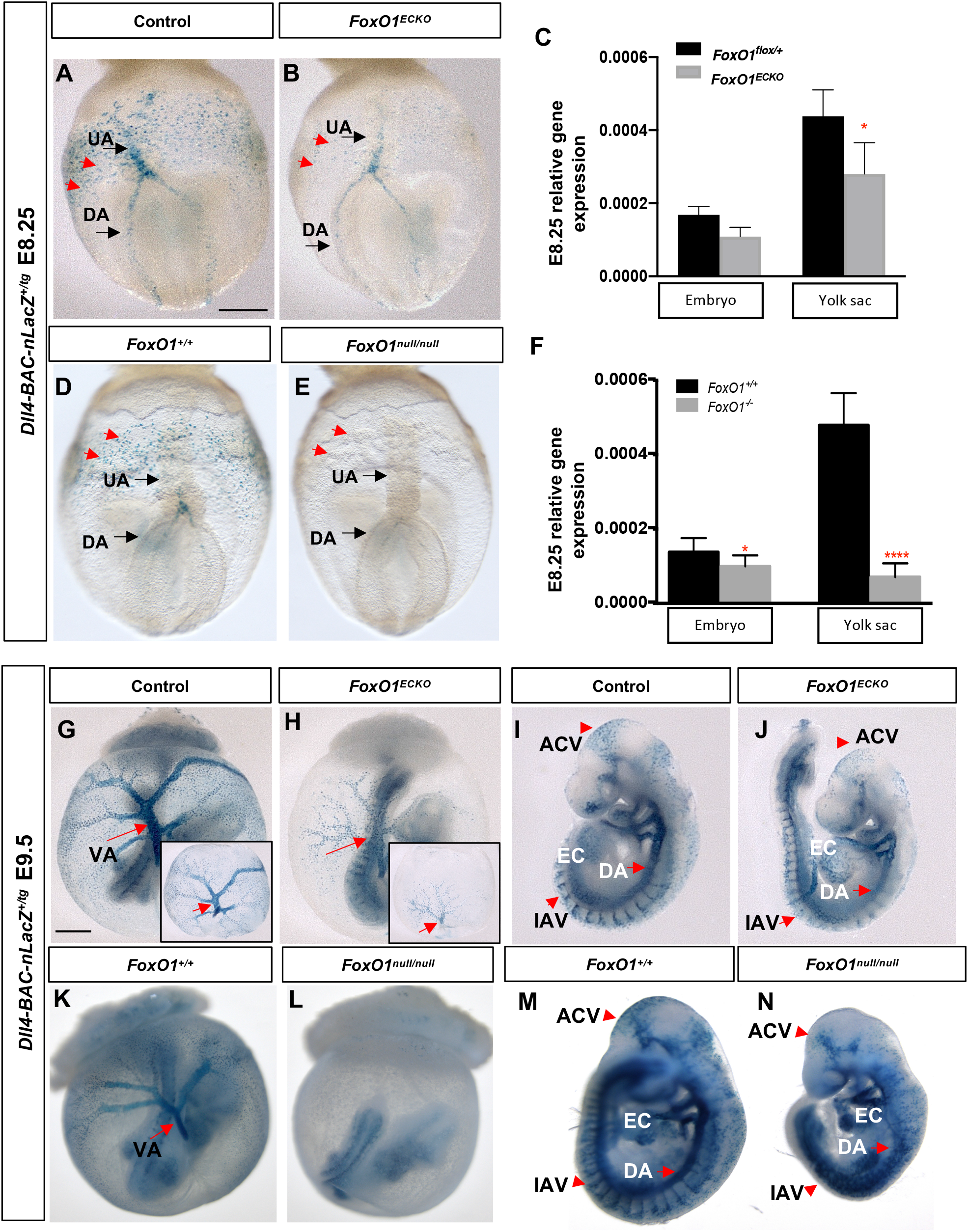
Characterization of arterial defects in *FoxO1*^*ECKO*^ and germline mutants using the *Dll4-BAC-nlacZ* reporter. (A and B, D and E) *nlacZ* reporter activity was detected in the dorsal aorta [DA] and umbilical artery [UA] in E8.25 control, *FoxO1*^*ECKO*^, and *FoxO1* null embryos. Arrowheads point to yolk sac endothelial cells in posterior region of yolk sac plexus. (C and F) *Dll4* expression in control, *FoxO1*^*ECKO*^, and *FoxO1* null embryos and yolk sacs. n>3. \**p*<0.05. (G and H, K and L) *nlacZ* reporter activity in E9.5 control, *FoxO1*^*ECKO*^, and *FoxO1* null yolk sac and embryo; VA = vitelline artery [arrow]; insets in G and H show yolk sacs only. (I and J, M and N) *nlacZ* reporter activity in E9.5 control, *FoxO1*^*ECKO*^, and *FoxO1* null; EC = endocardium, DA = dorsal aorta, IAV = intersomitic arterial vessels, ACV = arterial cranial vasculature. Scale bars for E8.25 panes = 200μm; E9.25 = 500μm.

By E9.5, nLacZ reporter expression was detected in the arterial tree in the yolk sac, particularly in the VA and within the arterioles (Figure 5G and 5G inset). Consistent with the E8.25 results, we found Dll4 expression reduced in E9.5 *FoxO1*^*ECKO*^ yolk sacs (Figure 5H and 5H inset) but strong expression in vessels within control and *FoxO1*^*ECKO*^ embryos (Figures 5I and J). Germline null embryos showed similar nLacZ activity compared to controls (Figure 5K), but reporter expression was not detectable in null yolk sacs, despite the strong expression seen in the embryo (5L). Isolated embryos confirmed LacZ expression in both control (5M) and null embryos (5N). Additional analysis of *Dll4* reporter expression confirmed previous reports of vascular defects in *FoxO1* mutants, including within the intersomitic vessels, cranial vessels and dorsal aorta (5I, J,M, and N, red arrows) (Hosaka et al., 2004, Ferdous et al., 2011, Dharaneeswaran et al., 2014). Collectively, our results indicate that FOXO1 plays a critical and early role in the regulation of *Dll4* expression, a key factor in determining arterial identity, within the extra-embryonic arteries of the yolk sac, but does not appear to be required for *Dll4* expression within the embryo proper.

### *FoxO1* deletion in endothelial cells upregulates *Sprouty2*/*4* expression

We uncovered a novel role for FOXO1 in the establishment of arterial identity, but neither *Flk1* nor *Dll4* contain known binding sites for FOXO1, so we examined expression levels of previously validated FOXO1 direct transcriptional targets in the endothelium: adrenomedullin (*Adm)*, BMP binding endothelial regulator (*Bmper*), *eNOS, Sprouty2*, and *Vcam1* (Potente et al., 2005, Ferdous et al., 2011). qRT-PCR analysis in *FoxO1*^*ECKO*^ yolk sacs at E8.25 (Figure 6A) revealed no significant reduction in *Adm* or *Bmper*, but a significant downregulation in *eNOS* and *Vcam1*, as previously described in other tissues (Potente et al., 2005, Ferdous et al., 2011). Interestingly, *FoxO1*^*ECKO*^ yolk sacs showed significantly increased *Sprouty2* expression (Figure 6A). Subsequent analysis showed that in addition to *Sprouty2, Sprouty4* was also upregulated in mutant yolk sacs compared to controls, while *Sprouty1* and *3* levels were unchanged (Figure 6B). We focused our subsequent analysis on the Sprouty factors because Sprouty4 over-expression had been shown to inhibit angiogenesis in the yolk sac (Lee et al., 2001) and the upregulation of *Sprouty* transcripts led us to hypothesize that FOXO1 may act as a direct repressor of *Sprouty* gene expression.

**Figure 6.**
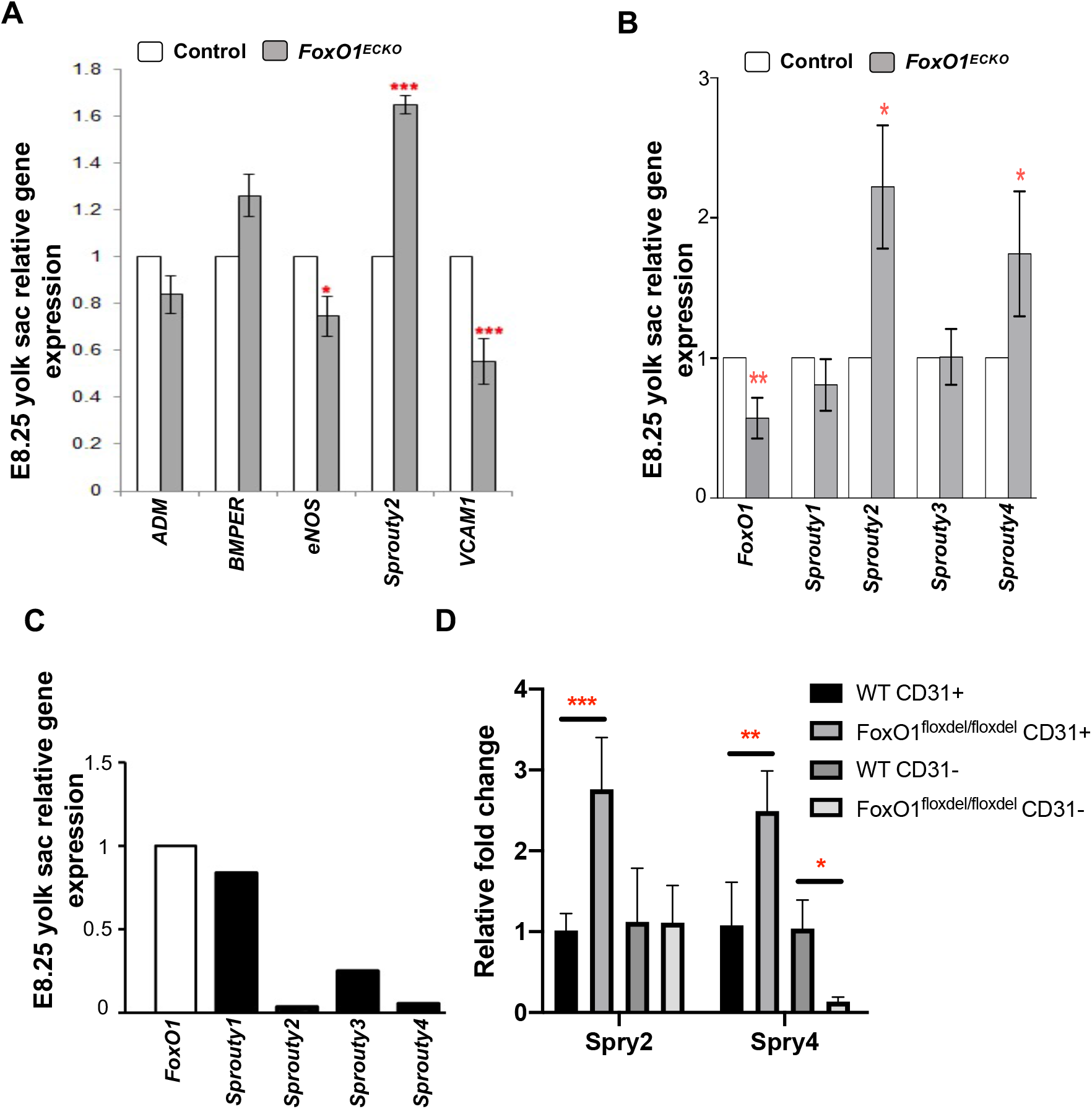
FOXO1 regulates *Sprouty2*/*4* expression in the yolk sac vasculature. Quantitative RT-PCR analysis in E8.25 control and *FoxO1*^*ECKO*^ yolk sacs for (A) known FOXO1 targets and (B) *Sprouty* family members. (C) Quantitative RT-PCR of endogenous *Sprouty1-4* expression relative to *FoxO1*. (D) Quantitative RT-PCR of *Sprouty2/4* in MACS sorted E8.25 CD31+ and CD31-control and *FoxO1* null yolk sac cells.

In keeping with the idea that FOXO1 may normally repress *Sprouty 2/4* transcription, we examined endogenous mRNA levels of *FoxO1* and *Sprouty1-4* in wildtype E8.25-8.5 yolk sacs. This analysis revealed that *Sprouty2/4* expression is much lower than *FoxO1, Sprouty1* or *Sprouty3* (Figure 6C). To expand and confirm our previous analysis of *Sprouty 2/4* expression, we examined *Sprouty 2/4* transcripts in CD31^+^ and CD31^-^ MACS-sorted ECs from germline *FoxO1* mutants and controls (Figure 3D). For *Sprouty2*, we observed increased expression in CD31^+^ cells, but there was no change in non-endothelial cells from null mutant yolk sacs (Figure 6D). Surprisingly, while *Sprouty4* transcripts increased in ECs of null yolk sacs, transcripts were decreased in non-endothelial cells. These data suggest that FOXO1 may act as both a transcriptional repressor or activator in adjacent tissues in the yolk sac, depending on the cell identity or transcriptional target.

### FOXO1 directly binds to endogenous *Sprouty2/4* promoters and represses *Sprouty2/4* transcription

FOXO1 is known to regulate *Sprouty2* mRNA expression in liver endothelial cells, and *in vivo* chromatin immunoprecipitation (ChIP) experiments confirmed that FOXO1 occupies four conserved FOXO binding elements within the murine *Sprouty2* locus (Paik et al., 2007). The first FOXO1 binding site (Figure S5A) is located ∼4kb upstream of the transcriptional start site (TSS) of murine *Sprouty2*, and the second DNA-binding site is within exon 2 (Figure 7A). The third and fourth FOXO1 binding sites are located ∼5kb and 7kb downstream of the TSS, respectively (Figure 7A). Paik, et al. showed that FOXO1 interacts with these loci to activate *Sprouty2* in the liver, but it was unknown whether FOXO1 utilizes the same binding sites to repress *Sprouty2* in the yolk sac. To determine if FOXO1 occupies any of these four identified binding sites in the murine yolk sac, we performed ChIP-PCR using pooled E8.25 yolk sacs. As shown in Figure 7B, FOXO1 occupancy was significantly enriched at the -4051, +5060, and +6972 regions compared to IgG control. FOXO1 enrichment was not observed at the +4479 region. This demonstrates that during early yolk sac blood vessel development, FOXO1 binds *Sprouty2* at regulatory regions -4051, +5060, and +6972 and supports the context-dependent function of FOXO1.

**Figure 7.**
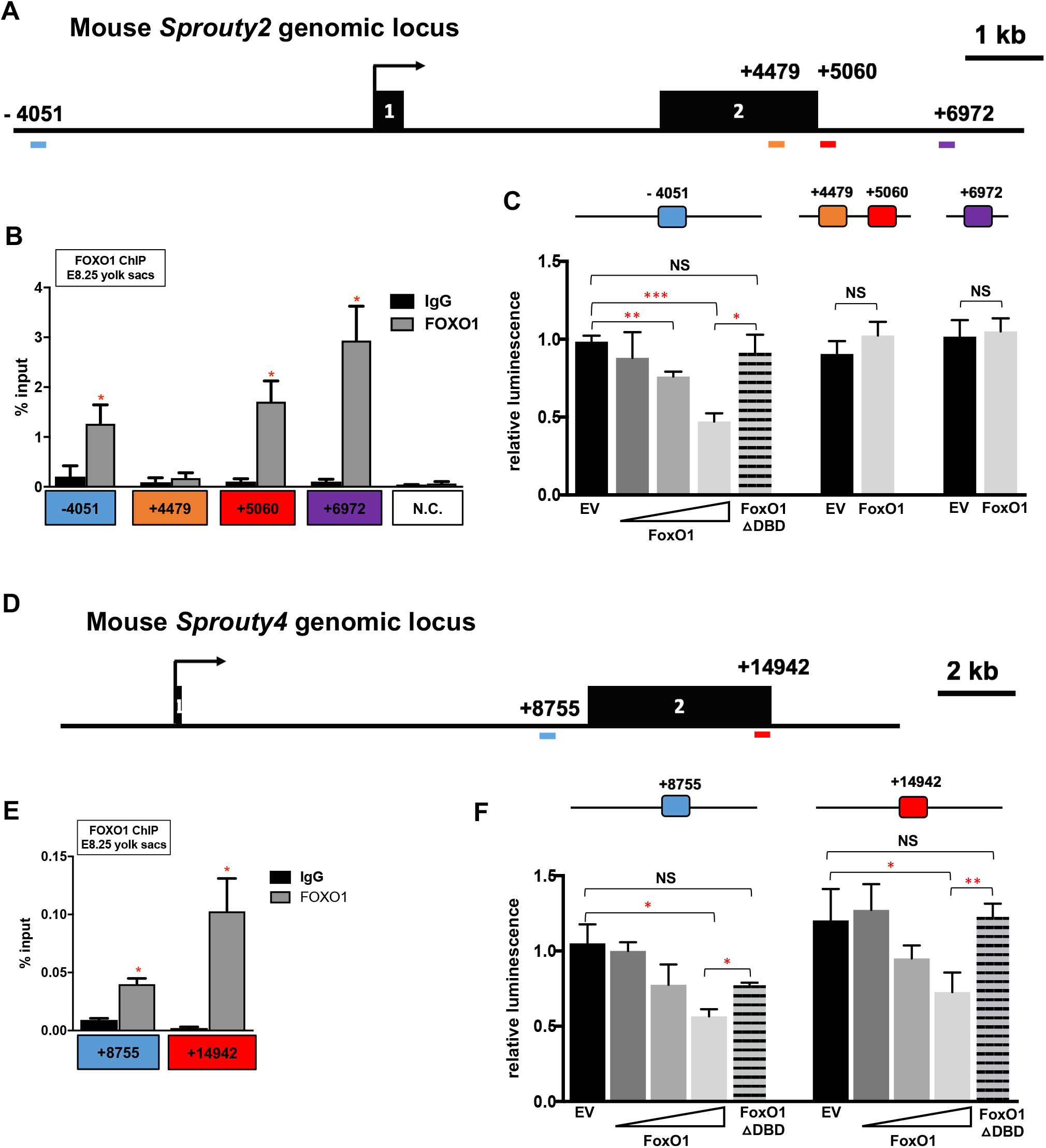
FOXO1 directly binds to endogenous *Sprouty2/4* promoters and represses *Sprouty2/4* transcription. (A) Genomic locus of mouse *Sprouty2* gene with FOXO1 binding sites in red. (B and E) FOXO1 ChIP-PCR using E8.25 yolk sac chromatin. (C) Luciferase activity of FOXO1 on *Sprouty2* promoter in H1299 cells. (D) Genomic locus of mouse *Sprouty4* gene with FOXO1 binding sites in red. (F) Luciferase activity of FOXO1 on *Sprouty4* promoter in H1299 cells. \**p*<0.05, \*\**p*<0.01, \*\*\**p*<0.001. EV, empty vector.

Next, we generated luciferase reporter constructs containing ∼2kb of the murine *Sprouty2* promoter, or specific regulatory regions harboring FOXO1 binding sites, and measured transcriptional activity in cultured mammalian cells (Figure S5B). To avoid potential confounds in our analysis from endogenous FOXO1, the human lung cancer cell line H1299 was chosen since FOXO1 protein expression is undetectable in this line (Zhao et al., 2010). Overexpression of *FoxO1* (*FLAG::FoxO1*) significantly repressed luciferase activity of the promoter construct containing the -4051 FOXO1 binding site in a dose dependent manner. Furthermore, co-transfecting the same reporter construct along with a FOXO1 cDNA without a DNA binding domain abolished this transcriptional repression (Figure 7C). In contrast, FOXO1 did not significantly repress luciferase activity in the constructs containing either the +4479/5060 or +6972 *Sprouty2* regulatory regions. These results suggest that FOXO1 directly downregulates *Sprouty2* expression via the -4051 site in its promoter.

To determine if this role for FOXO1 is evolutionarily conserved, we examined the *Sprouty4* locus for conserved FOXO1 DNA-binding motifs. Two putative binding sites, which were conserved in at least three vertebrate genomes (mammalian and non-mammalian), were identified +8755 bp and +14942 bp downstream of the *Sprouty4* TSS (Figure 7D). FOXO1 ChIP-PCR using E8.25 yolk sac chromatin showed a significant enrichment of FOXO1 occupancy in both regulatory regions (Figure 7E). Luciferase assays in H1299 cells also showed that these same sites were required for wildtype FOXO1 dose-dependent repression of reporter activity (Figure 5F; S5E). Taken together, data from the *Sprouty* mRNA expression analysis, as well as ChIP and luciferase assays, demonstrated that FOXO1 directly repressed *Sprouty2* and *Sprouty4* transcription in the E8.25 murine yolk sac via known and newly identified conserved DNA-binding sites.

### Transient overexpression of *Sprouty4* in endothelial cells phenocopies conditional loss-of-function *FoxO1* mutants

Given the fact that *Sprouty2* and *Sprouty4* are known to have anti-angiogenic functions (Taniguchi et al., 2009, Wietecha et al., 2011, Lee et al., 2001), and our data herein show that FOXO1 directly represses *Sprouty2/4* expression in the yolk sac, we hypothesized that FOXO1 promotes arterial gene expression by repressing *Sprouty2/4*. It had previously been shown that adenovirus-mediated overexpression of *Sprouty4* in developing embryos inhibited sprouting and branching of small vessels in the embryo proper and vessel remodeling in the yolk sac (Lee et al., 2001). To test whether *Sprouty4* overexpression could recapitulate the *FoxO1* loss-of-function phenotype, we utilized a well-characterized *Flk1* promoter-enhancer construct (Kappel et al., 1999, Ronicke et al., 1996, Fraser et al., 2005) to transiently overexpress *Sprouty4* in the endothelial cells of the mouse embryo and YS beginning at E7.5. To track transgene expression, a H2B::YFP reporter was inserted downstream to enable identification of YFP^+^ transgenic embryos (schematized in Figure 8A). E8.25 or E9.5 yolk sacs of YFP^+^ transgenic embryos (n=3) showed poorly remodeled vasculature, as their vessels remained as a primitive vascular plexus, while at E9.5 non-transgenic embryos had a normally developed vitelline artery with large caliber vessels branching into smaller diameter capillaries (Figure 8B). The lack of yolk sac vascular remodeling in the transient transgenic *Flk1*-*Sprouty4* embryos phenocopied the *FoxO1*^*ECKO*^ embryos and was similar to previous loss of function data (Lee et al., 2001). Yolk sacs harvested from both transgenic and control embryos confirmed the expression of exogenous *Sprouty4* and detection of YFP transcripts only in transgenic embryos (Figure S6).

**Figure 8.**
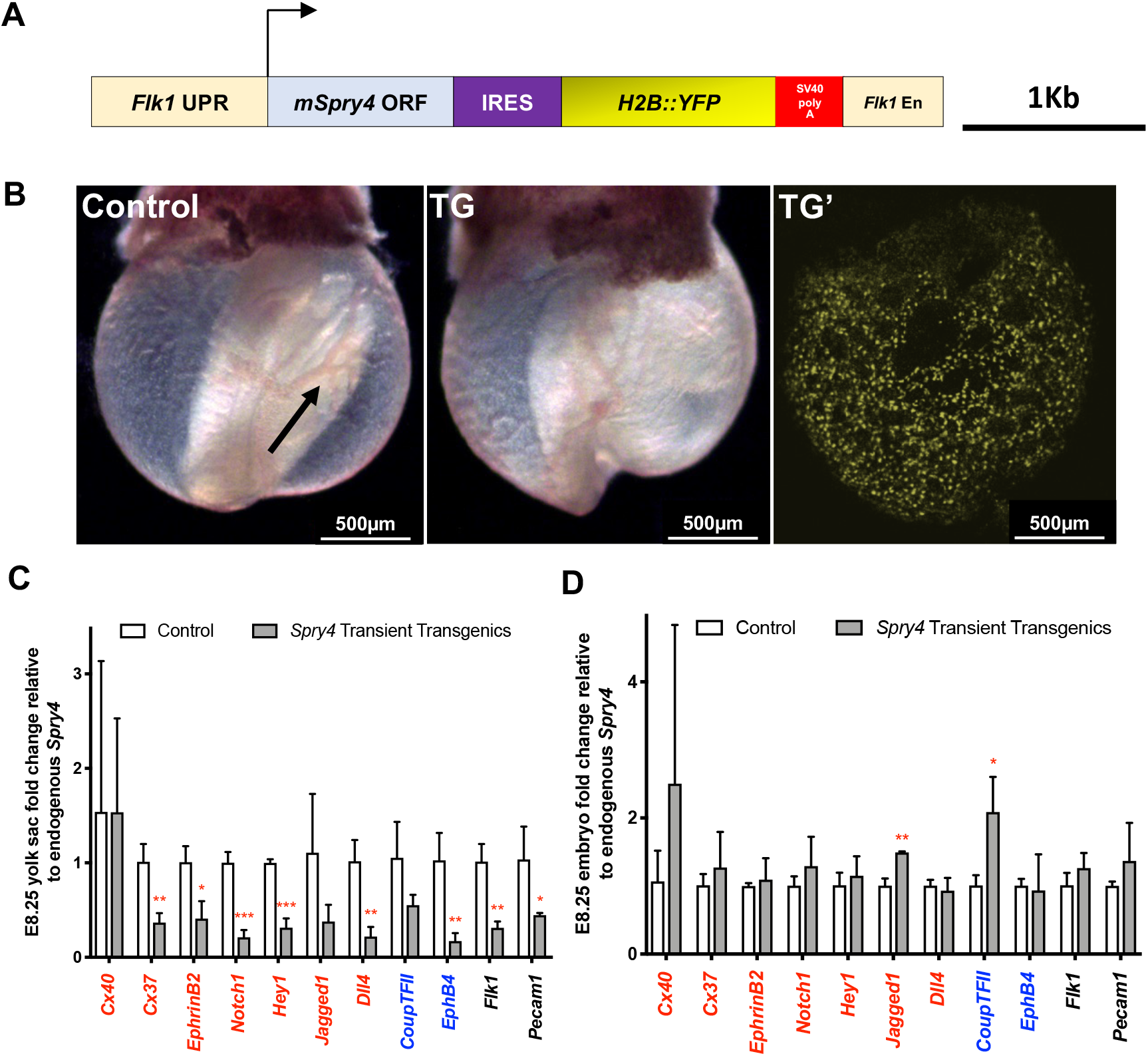
Transient overexpression of *Sprouty4* in endothelial cells phenocopies conditional loss-of-function *FoxO1* mutants. (A) Schematic of Sprouty4 overexpression construct for pro-nuclei injection. (B) Brightfield image of E9.5 non-transgenic (control) and transgenic embryo (TG); confocal imaging of TG embryo showing YFP fluorescence in yolk sac. Note vessel remodeling in the control yolk sac (arrow). Quantitative RT-PCR of arterial markers in control and TG yolk sacs (C) and embryos (D) (n=3). \**p*<0.05, \*\**p*<0.01, \*\*\**p*<0.001.

Next, we collected total RNA from E8.25 transgenic YFP^+^ and control yolk sacs and embryos for analysis of arterial marker genes. The relative expression level of each arterial marker was normalized to relative endogenous *Sprouty4* in order to compare the effect of exogenous *Sprouty4* overexpression, and then compared between control and YFP^+^ transgenic groups. Arterial markers, such as *Cx37, EphrinB2, Notch1, Hey1, Jagged1*, and *Dll4* were significantly downregulated in the yolk sacs of transgenic embryos compared to controls (Figure 8C), while expression within the embryo proper of these markers was not significantly changed, with the exception of *Jagged1* (Figure 8D). The expression of *Cx40* was not significantly different between the control and transgenic groups in either the yolk sac or embryos (Figures 8C and D). Additionally, unlike in *FoxO1*^*ECKO*^ yolk sacs, expression of venous marker *EphB4* was significantly down regulated in the yolk sac and *Coup-TFII* expression was significantly increased in the transgenic embryos, suggesting that either Sprouty4 could have FOXO1 independent functions, or that abnormally high levels of *Sprouty4* may affect other processes. These data, combined with our results showing that FOXO1 represses *Sprouty2/4* transcription, indicates that FOXO1 acts as a key transcriptional regulator in arterial-venous specification by repressing an antagonist of arterial specification.

## DISCUSSION

Previously, using either through germline mutations or conditional approaches, several groups demonstrated a requirement for FOXO1 in the early embryo, as these mutants featured failed yolk sac remodeling and mid-gestation lethality (Sengupta et al., 2012, Furuyama et al., 2004, Hosaka et al., 2004). In this paper, we investigated the role of FOXO1 within endothelial cells prior to the onset of consistent circulation and overt vascular remodeling. It is well known that arteriovenous specification and the arterial gene expression program are influenced by hemodynamic forces (le Noble et al., 2004, le Noble et al., 2005, Wragg et al., 2014), but our goal here was to determine whether FOXO1 functions in the endothelium before the onset of hemodynamic signaling to affect arteriovenous patterning. Herein, we demonstrate that blood flow is normal in *FoxO1*^*ECKO*^ mutants at these early stages, although heart failure and poor circulation are evident by E9.5 (Figure 2). Others have shown that FOXO1 is not required for heart development (Sengupta et al., 2012), and it is possible that heart failure in these embryos is caused by the increased resistance of blood flow encountered in the unremodeled vitelline vessels. It has also been noted that loss of FOXO1 causes allantois defects, preventing normal allantois fusion and circulation to the placenta (Ferdous et al., 2011). We did not observe overt defects in allantois fusion in *FoxO1*^*ECKO*^ embryos (0/10 *FoxO1*^*flox/flox*^*;Tie2-cre*^*Tg/+*^) (Table 1) and observed only low penetrance of allantois fusion defects in (2/15) in the germline *FoxO1* knock out embryos, whereas 100% of all null or *FoxO1*^*ECKO*^ embryos examined showed defects in yolk sac remodeling, heart failure and mid-gestation lethality. Thus, it is likely that the heart failure and lethality are caused by increased resistance to blood flow in vitelline vessels; however, the reduction in *Dll4* expression that we observed using the *Dll4-BAC-nlacZ* reporter in *FoxO1*^ECKO^ and germline null mutants suggest that further investigation of the consequence *Dll4* loss in allantois development is warranted.

In this study, we report that FOXO1 plays a previously unidentified role in regulating arterial-specific gene expression prior to the onset of blood flow. Based on transcript expression analyses, antibody immunostaining, and transgenic murine reporter experiments, we have concluded that loss of *FoxO1* causes a significant downregulation in *Flk1* and other critical arterial markers, including *Dll4*, in YS endothelial cells without affecting cell proliferation or cell death. Further, our data demonstrate that FOXO1 directly binds to regulatory regions of *Sprouty 2* and *4* in the yolk sac, and that FOXO1 acts as a direct repressor of *Sprouty 2* and *4* in endothelial cells. Finally, we show that the overexpression of *Sprouty4* in endothelial cells *in vivo* was sufficient to recapitulate impairments in both vascular remodeling and arterial cell fate specification seen in *FoxO1* mutants. Thus, these data indicate that repression of *Sprouty2/4* by FOXO1 is required to promote early, specification of arterial identity in the yolk sac prior to the onset of robust embryonic circulation.

Interestingly, although we observed elevated *Sprouty 2/4* transcripts in yolk sac endothelial cells of *FoxO1* null embryos, we found reduced *Sprouty 4* mRNA expression in non-endothelial cells (CD31-) in the yolk sac, suggesting FOXO1 may act as activator for *Sprouty 4* in other cell types of the yolk sac. Several recent studies have shown that FOXO1 functions as a transcriptional repressor in hepatocytes (Langlet et al., 2017) and pancreatic progenitor cells (Jiang et al., 2017). Interestingly, in some instances co-factors have been identified that enable FOXO1 to act as an activator or a repressor within the same tissues (Langlet et al., 2017). A similar mechanism may explain the observed differences between endothelial and non-endothelial cells within the yolk sac. It is not yet known whether a transcriptional co-factor in YS endothelial cells is required for FOXO1 to act as a repressor, or if another mechanism accounts for the opposite regulation of *Sprouty 4* in adjacent cell layers.

*FoxO1* loss did not appear to effect normal expression of pan-endothelial markers *Pecam1, Tie2*, and other genes, but *Flk1/Vegfr2* was significantly reduced in both *FoxO1*^*ECKO*^ and sorted germline mutant yolk sac endothelial cells. We also found that despite the reduction in *Flk1/Vegfr2*, we did not observe changes in cell proliferation or cell viability, two process that are directly regulated by VEGF-VEGFR signaling (Bernatchez et al., 1999). Sprouty factors inhibit receptor tyrosine kinase (RTK) signaling, and Sprouty overexpression could cause a reduction in *Flk1/Vegfr2* that is normally promoted by FLK1 or FGF receptor activation (Lee et al., 2001, Casci et al., 1999). Indeed, we observed a reduction in *Flk1* transcripts when *Sprouty4* was overexpressed in transgenic embryos, but also observed a strong effect on the expression of other endothelial markers such as *Pecam*. It is also possible that the reduction of *Flk1* expression seen in *FoxO1* mutants is a secondary consequence of disrupted arterial specification, rather than a primary driver of this defect.

*Dll4* is among the earliest markers of arterial gene expression (Chong et al., 2011, Wythe et al., 2013), but precisely how *Dll4* expression is initiated within the early yolk sac and embryo remains poorly understood. While *Dll4* transcription was suggested to be regulated by 5’ binding of FOXC1/2 and β-catenin in its proximal promoter (Corada et al., 2010, Hayashi and Kume, 2008, Seo et al., 2006), subsequent *in vivo* analysis showed that this region is not sufficient to mediate expression (Wythe et al., 2013). Additionally, endothelial-specific loss of *β-catenin* failed to alter *Dll4* expression in mice, or produce arteriovenous patterning defects (Wythe et al., 2013). Furthermore, functional enhancers were found within intron 3 and upstream at -12 and -16 kb that recapitulated the pattern of endogenous *Dll4* expression (Sacilotto et al., 2013, Wythe et al., 2013), and these regions lacked conserved FOXC1/C2 binding sites, as well as TCF/LEF binding sites. In these current studies, we used a nuclear localized LacZ reporter that recapitulates the normal expression pattern of *Dll4* (Herman et al., 2018). Our data clearly showed that the reduction in *Dll4* expression was far more severe in the yolk sac of *FoxO1*^*ECKO*^ or null embryos than within the embryo proper. Similarly, the overexpression of *Sprouty 4* throughout the embryo and yolk sac using an endothelial-specific *Vegfr2/Flk1* promoter (Kappel et al., 1999, Ronicke et al., 1996, Fraser et al., 2005) indicated that *Sprouty* overexpression did not alter arterial gene expression in the embryo, but suppressed arterial transcripts in the yolk sac. The mechanism that explains the differential activity of FOXO1 and SPROUTY2/4 within the vasculature of the yolk sac vs the embryo proper remains unclear. Future experiments will be required to address numerous possible explanations including differences in mesodermal cell lineages, differential binding to cofactors and/or differences in post-translational modifications regulated by local cell-cell signaling.

One unresolved question from these studies is the relationship between abnormal arteriovenous specification and failed vessel remodeling. Both arteriovenous identity and vessel remodeling are regulated by hemodynamic forces, and AV specification relies on pathways that respond to VEGF signaling (Fish and Wythe, 2015, Covassin et al., 2006, Weinstein and Lawson, 2002, Fang et al., 2017). In *FoxO1*^*ECKO*^, we detected a downregulation in Flk1/VEGFR2. VEGFR2 and other VEGF receptors have been shown to act as shear stress mechanosensors, signaling through downstream pathways such as the MEK-ERK kinase cascade in response to changes in blood flow (Tzima et al., 2005, Baeyens and Schwartz, 2016). Thus, the downregulation of VEGFR2 in *FoxO1* mutant embryos could prevent ECs from responding to normal blood flow signaling needed for vessel remodeling. Previously, our lab showed that ECs within the vitelline arteries, but not the vitelline veins, migrate directionally in response to hemodynamic changes in the yolk sac vasculature (Udan et al., 2013) so it is possible that the loss of FOXO1 and/or the overexpression of *Sprouty2/4* interferes not only with initial *Dll4* specification, but with the cell’s ability to sense mechanical signaling that is necessary to direct cell migration required for remodeling. Further work will be needed to better understand the mechanisms leading both to early *Dll4* expression and those that regulate the cellular responses needed for vessels to adapt to changes in blood flow.

## EXPERIMENTAL PROCEDURES

### Animals and genotyping

All animal experiments were conducted according to protocols approved by the Institutional Animal Care and Use Committee of Baylor College of Medicine. *Ella-Cre* and *Tie2-Cre* transgenic mice were purchased from Jackson Labs (# 003724 and 008863, respectively). *ɛ-globin-KGFP* (Dyer et al., 2001), *FoxO1*^*flox/flox*^ mice (Paik et al., 2007), and *Dll4-BAC-nLacZ* mice (Herman et al., 2018) were maintained and genotyped as previously described. *FoxO1* germline knockout mice were generated by crossing the *FoxO1*^*flox/flox*^ mice to *Ella-Cre* mice (Lakso et al., 1996). *Flk1-H2B::YFP* reporter mice were kindly provided by Dr. K. Hadjantonakis, Memorial Sloan Kettering Cancer Center (Fraser et al., 2005).

### Immunostaining of whole or sectioned yolk sacs and LacZ staining

E8.25 yolk sacs were fixed in 4% PFA, rinsed in PBS, permeabilized with 0.1% TritonX-100 for 1 hour, and blocked in 2% normal donkey serum/1% BSA for 5 hours. Yolk sacs were then incubated with anti-PECAM1 antibody (BD Pharmingen, #550274; 1:100) overnight at 4°C. After several PBS washes, yolk sacs were incubated with goat anti-rabbit antibody (Molecular Probes, AlexaFluor 633,1:500) and DAPI (1:500) overnight at 4°C. Finally, yolk sacs were rinsed in PBS and imaged using the Zeiss LSM510 META confocal microscope. Dissected yolk sacs were cryosectioned at 20μm and sections were permeabilized and blocked, and incubated with antibodies to either Caspase 3 (Cell Signaling #9661, 1:50); Connexin37 (ThermoFisher Scientific #404200, 1:50); Connexin40 (ThermoFisher Scientific #364900, 1:50); eNOS (Santa Cruz #sc-654, 1:50); Flk1 (Sigma #V1014, 1:100); or pHistone H3 (Millipore #06-570, 1:50) overnight at 4°C. The secondary antibody incubation and image acquisition were performed as described previously.

*Dll4-BAC-nLacZ* transgenic reporter were examined on *FoxO1* germline or *FoxO1*^*ECKO*^ (*Tie2-Cre*^*+/tg*^*;FoxO1*^*floxfloxl*^) backgrounds. E8.5 and E9.5 embryos were dissected in cold PBS and fixed in 4% PFA. Embryos were then washed in X-gal rinse buffer (0.02% NP40, 0.01% sodium deoxycholate; 4 × 15mins) and thereafter stained in x-gal solution [5mM K_3_Fe(CN)_6,_ 5mM K_4_Fe(CN)_6,_ 0.01% sodium deoxycholate, 0.02% NP40, 2mM MgCl_2,_ 5mM EGTA, 1mg/ml X-gal] at 37°C overnight. Embryos were then post-fixed in 4% PFA and then cleared in 50% and 70% glycerol. Embryos from the same litter were processed and stained in a 20ml scintillation vial. Stained embryos were photographed using the Axio ZoomV16 (Zeiss) stereo microscope and thereafter genotyped.

### Quantification of *Flk1-H2B::YFP*^*+*^ cell density, proliferation index and apoptotic index in whole mount yolk sacs using FARSIGHT

Acquired WT and ECKO yolk sac whole mount images (n>3 yolk sacs per genotype, n>3 regions of interest per yolk sac) were made into maximum intensity projections and separated into individual RGB images: Red (pHistone-H3/Caspase 3), Green (Flk1-H2B::YFP) and Blue (DAPI). Individual nuclei for Red, Green and Blue channels were segmented and quantified using FARSIGHT, courtesy of Badri Roysam, University of Houston, which makes use of both intensity and volume thresholds to distinguish two nuclei as separate. YFP^**+**^ cell density was defined as the ratio of YFP^**+**^ nuclei to DAPI^+^ nuclei within that same field of view. Proliferative/apoptotic index was defined as the ratio of PH3^**+**^/ Caspase3^**+**^ nuclei to the number of DAPI^+^ nuclei. Endothelial cell proliferative/apoptotic index was defined as the ratio of YFP^**+**^;PH3^**+**^/Caspase3^**+**^ double positive nuclei to the number of YFP^**+**^ nuclei. The ratios were then averaged over the various WT and ECKO yolk sac images.

### Live imaging and analysis of blood flow in *FoxO1* conditional knockout embryos

*ε-globin-KGFP* reporter expressing GFP in primitive erythroblasts was examined in control and ECKO background and litters were dissected at E8.5 or E9.5 for blood velocity analysis. Embryos were dissected under a heated (37°C) dissection stage with warm dissection media (DMEM/F-12, 10% FBS, 100 U/mL penicillin, and 100 μg/mL streptomycin). Embryos with intact ectoplacental cone were placed in a glass bottomed culture chamber with culture medium (1:1 DMEM/F-12: rat serum, 100 U/mL penicillin, and 100 μg/mL streptomycin) and allowed to recover in a 37°C incubator for 20 minutes. Embryos were then placed on a heated confocal microscope stage (37°C) and imaged using the Zeiss LSM 5 LIVE laser scanning confocal microscope, using the Achroplan 20X/0.45 NA objective. A 200-frame time lapse (in a 512×512 pixel frame) was acquired at 30-50 frames per second. Blood flow time lapses were acquired at three different locations throughout the yolk sac per embryo, and at least three embryos of each genotype were used for data collection. Individual blood cell velocities in each track were determined from time lapse movies using Imaris. Individual blood cell velocities from three different locations per embryo were averaged. Average heart beats per minute were calculated by measuring the average time interval between peak velocities during the course of 5 cardiac cycles in individual velocity profiles for each embryo imaged. Embryos were genotyped after imaging, and blood velocities and heart beats per minute were averaged within WT and ECKO.

### Magnetic activated cell sorting of yolk sack endothelial cells

To isolate E8.25 yolk sac endothelial cells, fresh yolk sacs were dissected in cold DMEM/F12 media without phenol red (ThermoFisher Scientific #21041025), individually placed in 100 μL of cold TrypsinLE (Fisher Scientific #12605010), and kept on ice until all yolk sacs were harvested. Embryos were used for genotyping. To dissociate yolk sacs into single cell suspension, gently triturate with p200 pipette and incubate on ice for 5 minutes and repeat for a total of four times. To inhibit the enzyme, add 1 mL of stop solution: media + 10% FBS (ThermoFisher Scientific #26140079). The yolk sac cell suspensions were pelleted at 0.8 × 1000g for 5 minutes at 4 degrees Celsius. The pellets were resuspended in 90 μL of cell suspension buffer (PBS + 2% FBS + 2 mM EDTA). 10μL of CD31 MicroBeads (Miltenyi Biotec #130-097-418) were added to each yolk sac cell suspension and samples were incubated on ice for 15 minutes in the dark. Cell mixtures were pelleted at 0.8 × 1000g for 5 minutes at 4 degrees Celsius and washed with 1mL of cell suspension buffer. Cell mixtures were once again pelleted at 0.8 × 1000g for 5 minutes at 4 degrees Celsius and resuspended in 200uL of cell suspension buffer. Cell mixtures were passed through 40 μm cell strainers (Fisher Scientific #352340) into FACS tubes and strainers were washed with 300 μL of cell suspension buffer. MS columns (Miltenyi Biotec #130-041-301) were placed on OctoMACS separator and prepared according to manufacturer instructions. Cell mixtures were individually passed through columns and the flow through was reapplied through columns to maximize endothelial cell retention (CD31^+^ population). Columns were washed three times with 500 μL cell suspension buffer and all flow through was collected (CD31^-^ population). Bound cells were released from the columns by removal from magnetic separator, and 1 mL of cell suspension buffer was applied to the columns and cells were flushed using plunger into a 1.5 mL Eppendorf tube. Collected cells were then pelleted at 0.8 × 1000g for 5 minutes at 4 degrees Celsius and resuspended in Trizol (Thermo Fisher 15596018). After genotyping, CD31^+^ and CD31^-^ populations from two yolk sacs were combined and processed for RNA isolation (QIAGEN RNeasy Micro Kit #74004), cDNA synthesis and qRT-qPCR as described below.

### RNA isolation and qRT-PCR analysis

Total RNA was isolated from pooled E8.25 yolk sacs dissected from either *Tie2-Cre*^*+/tg*^; *FoxO1*^*+/flox*^ or *Tie2-Cre*^*+/tg*^; *FoxO1*^*flox/flox*^ embryos. Purified RNA was reverse transcribed (ThermoFisher Scientific #11752-050) and gene expression analysis was performed using TaqMan real-time assays for *FoxO1, Adm, Bmper, Vcam1*, and a panel of endothelial, arterial, and venous markers (see Fig. 2D, E for gene list). The data were normalized to *Gapdh* (Pfaffl, 2001) and relative expression ratios between control and ECKO embryos were determined. Endogenous *FoxO1* and *Sprouty*1-4 expression from pooled E8.25 yolk sacs (CD1 strain) was also probed by TaqMan real-time assay, but expression was calculated as fold change relative to *FoxO1*, which was set to 1.

Endogenous *Dll4* expression was measured in either germline *FoxO1* knockouts or *FoxO1* ^ECKO^ embryos at E8.25. The allantois was used for genotyping and total RNA was extracted from individual embryos and yolk sacs (n≥3) and probed for *Dll4* expression via TaqMan assay and fold change of expression between controls and homozygous *FoxO1* ^ECKO^ mutants, and statistical analysis were performed as described above.

### Chromatin Immunoprecipitation (ChIP) and qPCR

To determine endogenous FOXO1 chromatin occupancy, E8.25 yolk sacs from CD1 embryos were used for chromatin extraction. Freshly dissected yolk sacs were dissociated in ice cold PBS with protease inhibitors and the tissue was then crosslinked with 1.5% formaldehyde, followed by incubation with 125 mM glycine, and washed with PBS. After centrifugation, the pellet was resuspended in cell lysis buffer (5 mM PIPES, pH 8; 85 mM KCl; 0.5% NP40). The samples were spun and the pellet was resuspended in nuclear lysis buffer (50 mM Tris-HCl, pH 8.1; 10 mM EDTA; 1% SDS) and then sonicated on ice using a Bioruptor (Diagenode) to obtain sheared chromatin ranging between 100–500 bp. ChIP was performed according to the instructions for Magna ChIP kit (Millipore #17-10085) using 5 μg of anti-FOXO1 antibody (Abcam #ab39670) or rabbit IgG (Millipore #12370). The cross links were then reversed, and the purified DNA was then analyzed by qPCR in technical triplicates using SYBR green master mix and the primers listed in Table S1 to measure the percentage of co-precipitating DNA relative to input (% input) in *Sprouy2* and *Sprouty4* genomic regions.

### Cloning of *mSprouty2/4* promoter constructs and Luciferase Assay

Genomic regions of ∼2kb in length of murine *Sprouty2* and *Sprouty4* were PCR amplified using primers listed in Table S1, and using BAC clones of C57BL6 genomic DNA as template DNA (CH29-611D15, CH29-100M12, respectively; CHORI BAC/PAC resources). PCR fragments were ligated into pCRII-TOPO vector, sequenced, and then subcloned into pGL3-Promoter vector (Promega). H1299 cells (ATCC #CRL-5803) were maintained in DMEM media supplemented with 10% FBS, 100 U/ml penicillin, and 100 μg/ml streptomycin. For transient transfections, 50,000 cells were plated (48-well plate) and after 24 hours, each well was co-transfected with 200 ng of *Sprouty2/Sprouty4* promoter construct, 10 ng of pRL-TK, and 125 ng expression plasmid (pcDNA3-FOXO::FLAG, Addgene #13507) using manufacturer’s recommendations for lipofectamine 3000. The total amount of expression plasmid transfected per well was kept constant with varying amounts of pcDNA3.1 vector. As a negative control, a *FoxO1* plasmid encoding a deleted DNA binding domain (amino acid 208-220) was used (Addgene #10694). After 24 hours, cells were lysed and analyzed for firefly and *Renilla* luciferase activities according to the procedure outlined in the Dual-Glo luciferase assay system (Promega #E2920). All luciferase assays were performed in triplicates and repeated at least three times. Student’s *t* -test was used to assess statistical significance (*P* < 0.05 was considered statistically significant) and the averages and standard deviation from triplicate samples from representative assays were shown.

### Transient endothelial-specific *Sprouty* expression in embryos

A mouse *Sprouty4* cDNA clone (TransOmics clone BC057005) was used as a template to PCR amplify the coding sequence with 5’ *Sac*I and 3’ *Pme*I restriction sites (5’ GAGCTCCCAGCCTCA TGGAGCCC 3’ and 5’ GTTTAAACTCAGAAAGGCTTGTCAGAC 3’) and subcloned into pCRIITOPO vector (S4-2). The internal ribosome entry site (IRES) sequence was amplified from pIRES-hrGFP1a vector (Agilent Technologies) with 5’ *EcoR*V and 3’ *EcoR*I restriction sites using primers 5’ CTATAGATATCACCCCCCTCTCCCTA 3’ and 5’ GCATGAATTCGGTTGTGGCCATT ATCATCGTG 3’ and subcloned into pCRIITOPO vector (IRES-4). To assemble the final transgenic construct, clones S4-2 and IRES-4 were excised with 5’ *Ecl136*II and 3’ *Not*I, and 5’ *Not*I and 3’ *Ecl136*II, respectively, and co-ligated into *Flk1*-H2B/EYFP vector (kindly provided by Dr. K. Hadjantonakis, Memorial Sloan Kettering Cancer Center) via a blunt-ended *Hind*III site. The *Flk1*-H2B/EYFP vector has a well characterized *Flk1* promoter and intronic enhancer sequences which drive YFP expression in endothelial cells (Fraser et al., 2005). All clones were verified by DNA sequencing and the final transgenic construct was excised with 5’ *Sal*I and 3’ *Xba*I to purify a 4.5kb fragment for pronuclear microinjection, which was performed at the BCM Genetically Engineered Mouse Core. Transient transgenic embryos were dissected with yolk sac intact and initially screened for YFP expression using confocal microscopy. Gross morphology of the embryo and yolk sac vasculature was examined at E9.5 while arterial marker analysis (*C37, Cx40, Dll4, EphB2, Hey1*) was performed at E8.25, at which point embryos and yolk sacs of YFP positive and negative samples (n=3 each) were lysed in Trizol for RNA extraction. Transgene-positive embryos were also confirmed via qRT-PCR for YFP expression (data not shown) and also quantitating the ratio of exogenous over endogenous *mSprouty4* expression using transcript-specific primers. Detection was via the Sybr-green or Taqman assay (for arterial markers) and fold change of expression between YFP-positive and negative samples, and statistical analysis were performed as previously described.

## FIGURE LEGENDS

**Table S1.**
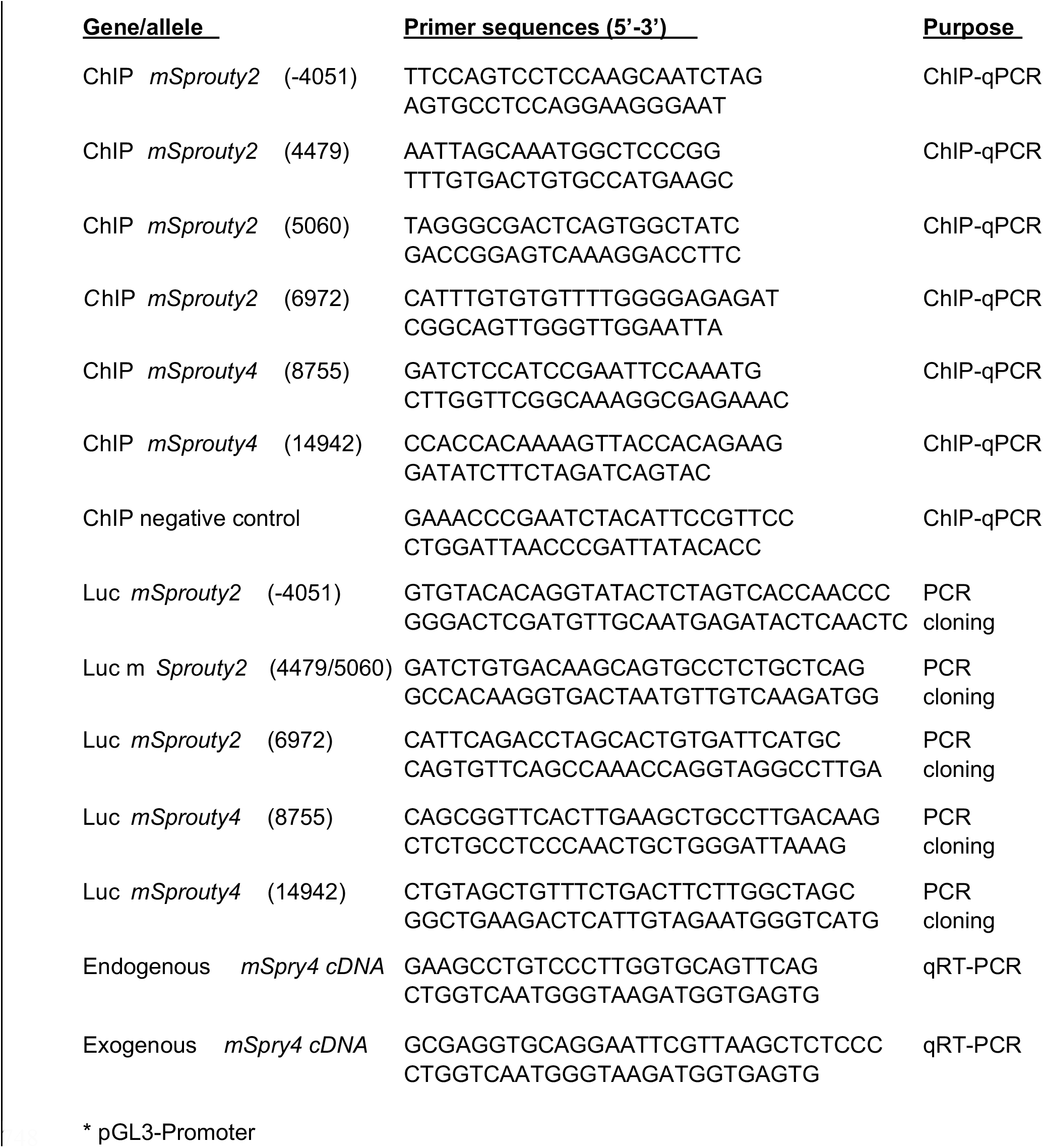
Primer sequences used for genotyping, ChIP-qPCR, and cloning

**Table S2.** Taqman assays for Gene expression analysis.

|  |  |  |  |
| --- | --- | --- | --- |
| *FoxO1* | Mm00490672_m1 | *Hey1* | Mm00468865_m1 |
| *FoxO3a* | Mm01185722_m1 | *Hey2* | Mm00468865_m1 |
| *FoxO4* | Mm00840140_g1 | *Jagged 1* | Mm00496902_m1 |
| *Flk1 (Kdr)* | Mm00840140_g1 | *Nrp1* | Mm00435379_m1 |
| *PECAM1* | Mm01242584_m1 | *Nrp2* | Mm00803099_m1 |
| *Tie2 (Tek)* | Mm01242584_m1 | *CoupTFII* | Mm00772789_m1 |
| *Flt1* | Mm00438980_m1 | *EphB4* | Mm01201157_m1 |
| *Connexin 43* | Mm00438980_m1 | *AFP* | Mm00431715_m1 |
| *eNOS* | Mm00435217_m1 | *ADM* | Mm00437438_g1 |
| *Connexin 37* | Mm00433610_s1 | *BMPER* | Mm01175806_m1 |
| *EphrinB2* | Mm01215897_m1 | *Sprouty2* | Mm00442344_m1 |
| *Notch1* | Mm00435249_m1 | *Sprouty4* | Mm00442345_m1 |
| *Dll4* | Mm00444619_m1 |  |  |

## Notes

### Competing Interest Statement

The authors have declared no competing interest.

